# SPgen: Proteome-wide Spatial Proteomics generation using multi-modality foundation models

**DOI:** 10.64898/2026.07.16.739037

**Authors:** Jiachen Li, Kaiyuan Yang, Qiaoling Che, Diwei Zheng, Wei Wei, Cheng Jin, Ye Yuan

## Abstract

Spatial proteomics (SP) measures the spatial distribution of proteins within tissues, providing important insights into tissue function, disease, and therapeutic response. However, current SP technologies profile only a small fraction of the proteome and are limited by cost and measurement noise. Recent AI approaches enable predicting spatial protein expression from transcriptomic or histopathological data, but are typically restricted to paired datasets covering only tens of proteins, limiting their ability to generalize beyond experimentally measured protein panels. Here we present SPgen, a multi-modal foundation-model framework for proteome-wide spatial protein prediction. SPgen integrates protein sequences, functional annotations, transcriptomic profiles, and spatial information to learn transferable representations that enable inference beyond experimentally profiled proteins. Across diverse spatial proteomics datasets, SPgen accurately reconstructs measured spatial patterns, reduces measurement noise, and enables proteome-wide spatial prediction.

## Introduction

Spatial proteomics (SP) enables direct measurement of protein distribution within tissues^1^, providing critical insights into development, immunity, tumor biology, and disease progression, etc^2,3^. Recent imaging- and mass spectrometry–based technologies have substantially expanded the scale of spatial protein profiling^4–6^. However, current SP platforms remain constrained by limited protein coverage, high cost^7^, and measurement noise. Even the largest SP datasets capture only a small fraction of the proteome, while important protein classes, including transcription factors (TFs) ^8^ and membrane proteins^9^, are often underrepresented owing to technical limitations. These challenges remain major barriers to comprehensive spatial characterization of the proteome.

Recent advances in artificial intelligence (AI) have enabled computational reconstruction and generation of SP data^7,10,11^. Existing approaches have enabled computational reconstruction of spatial proteomic data from transcriptomic or histopathological measurements^12–14^. However, these approaches are typically trained on paired datasets covering only tens of proteins and are therefore constrained by fixed protein panels defined by existing SP technologies, prohibiting unbiased proteome-wide spatial prediction.

Here, we developed SPgen, a multi-modal representation learning framework for proteome-wide spatial protein prediction. SPgen is built on the hypothesis that protein spatial organization is jointly determined by intrinsic protein properties and cellular context^15^. To model these complementary factors, SPgen jointly learns protein representations from protein sequences^16^, functional annotations^17^, and transcriptomic profiles^18^, and spot representations from SP data. By integrating protein and spot embeddings, SPgen learns transferable relationships between protein identity and spatial organization. As a result, SPgen could generate spatial distribution for proteins lacking direct measurements, extending inference beyond experimentally profiled panels. Across multiple mass spectrometry–based SP datasets, SPgen accurately reconstructs measured spatial patterns and generalizes to previously unseen proteins, including transcription factors and membrane-associated proteins.

## Results

### Overview of SPgen

SPgen aims to infer proteome-wide spatial protein distributions beyond the proteins measured in a given SP dataset. For this purpose, SPgen jointly learns protein representations and spatial-context representations to enable spatial prediction for unmeasured proteins (**Fig. 1**).

**Figure 1.**
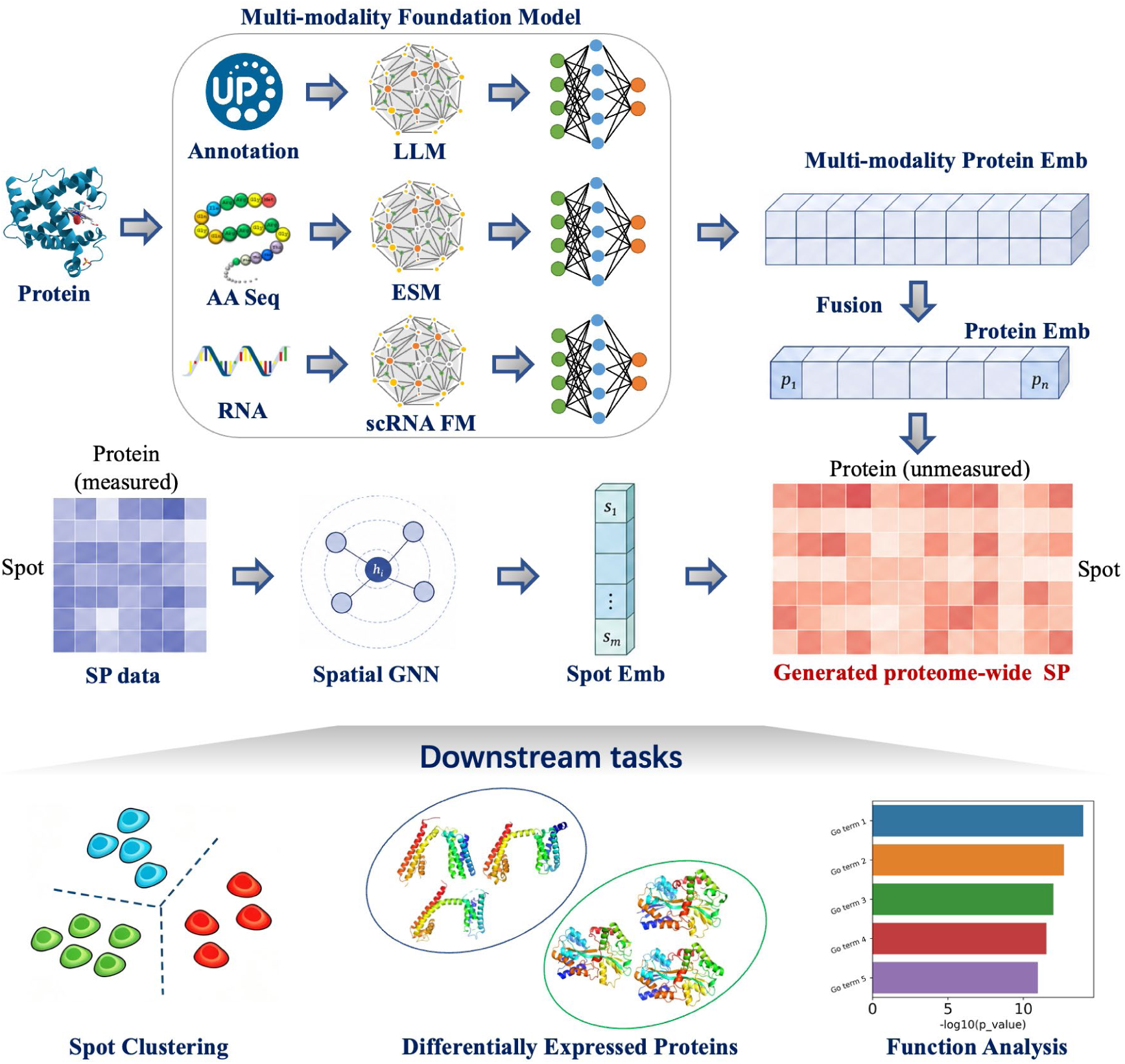
Overview of SPgen. SPgen jointly learns protein representations and spatial- context representations to enable spatial prediction for proteome-wide spatial prediction. Protein representations are learned using multi-modality foundation models that integrate embeddings from a LLM for annotation text, the ESM model for amino acid sequences, and a single-cell RNA foundation model (scRNA FM) for corresponding transcriptomic signals. Spatial-context representations are learned via a GNN constructed from the spatial distribution. SPgen generates proteome-wide SP data by integrating protein embeddings with spot embeddings. Downstream analyses are performed at both the protein level and the spot level.

Protein representations are derived from three complementary modalities: protein annotations, amino acid sequences, and gene-level transcriptomics. Specifically, SPgen encodes UniProt^17^ protein descriptions using a pretrained large language model (LLM)^19,20^ such as PubMedBERT^21^, protein sequences using a protein language model (for example, ESM^22^), and gene-level transcriptomics using pretrained single-cell foundation models^23,24^ (for example, scGPT^24^). These embeddings are integrated into a unified protein representation that captures functional, biochemical, and genomics profiles.

To model spatial context, SPgen learns spot representations from measured spatial proteomic profiles using a Graph Neural Network (GNN)^25,26^ that incorporates neighborhood information. During training, proteins are partitioned into training and testing sets, and only training proteins are used to construct spot representations, preventing information leakage. SPgen then predicts spatial abundance by combining protein and spot representations and is optimized (see **Methods** for details). Once trained, the model can infer spatial distributions for proteins not measured in the original experiment using only their pretrained protein representations.

### Proteome-wide SP generation on intestinal villus dataset by SPgen

We first evaluated SPgen on the intestinal villus spatial proteomics dataset generated by PLATO, comprising 2,093 spatial locations and 1,466 measured proteins. Proteins were randomly partitioned into training (90%) and testing (10%) sets. SPgen accurately reconstructed diverse spatial expression patterns for held-out proteins, with representative examples showing close agreement between predicted and measured distributions (**Fig. 2a**). Across testing proteins, the median Pearson correlation between predicted and observed abundances reached 0.74 (**Fig. 2b**).

**Figure 2.**
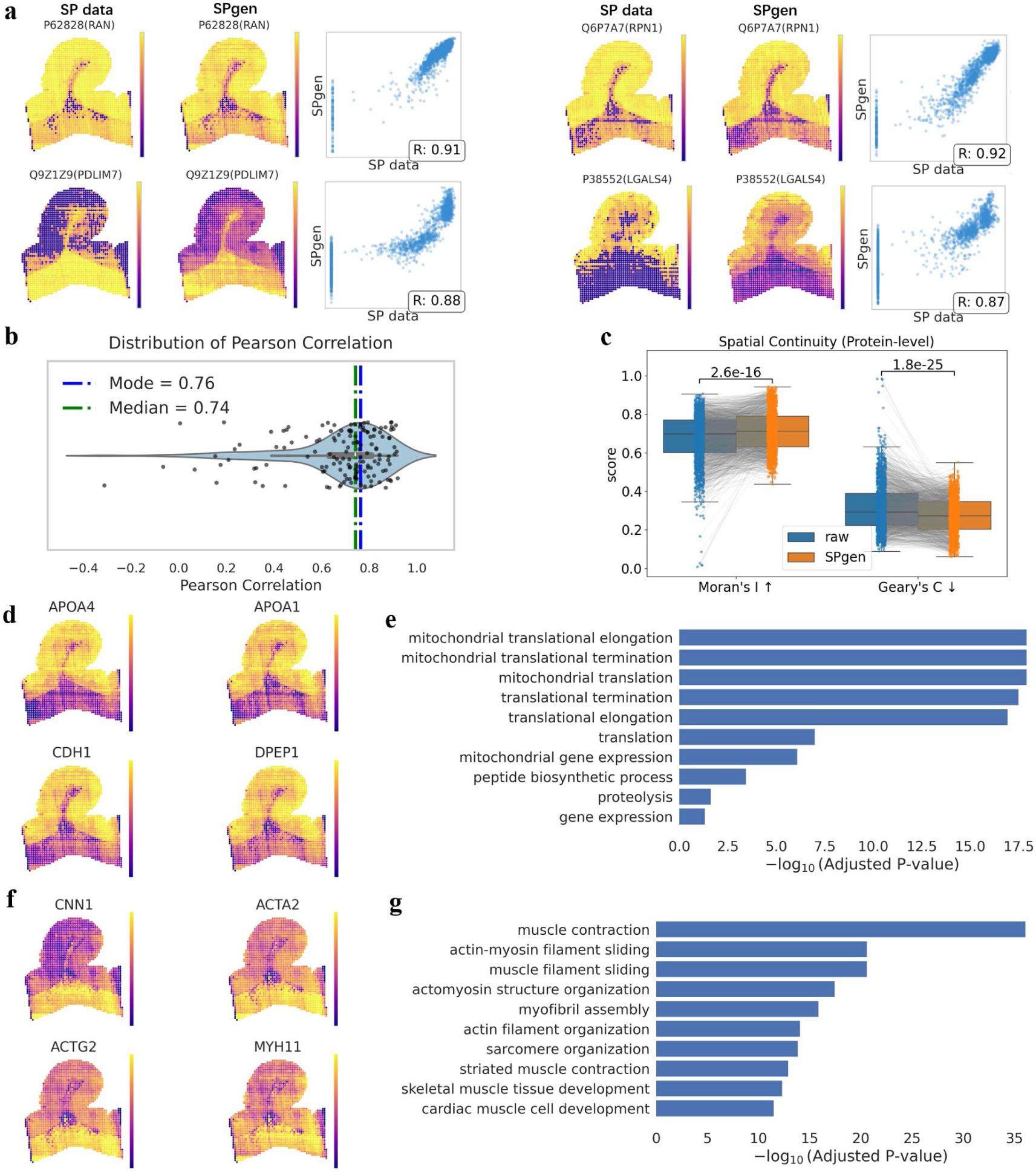
Results on the Intestinal villi SP dataset. (a) Representative examples of SPgen performance on the testing set, showing observed and predicted spatial distributions and their joint distribution. Pearson correlation coefficients are shown in the joint distribution panel. (b) Distribution of Pearson correlation on the testing set, with mode and median annotated. (c) Comparison of spatial continuity between SP data and SPgen predictions. Higher Moran’s I and lower Geary’s C indicate stronger spatial continuity. Paired samples are linked with gray lines. P-values of paired Wilcoxon tests are shown. (d,f) Predicted expression of representative proteins highly expressed in spot clusters 0 (panel d) and 1 (panel f). (e,g) GO enrichment analysis of the top 200 DE proteins in spot clusters 0 (panel e) and 1 (panel g).

In addition to recovering measured spatial patterns, SPgen generated spatial profiles with improved spatial coherence. Compared with the original measurements, SPgen predictions exhibited significantly higher Moran’s I and lower Geary’s C values, indicating stronger spatial autocorrelation and reduced local noise (**Fig. 2c**). These results suggest that SPgen not only reconstructs spatial proteomic signals but also effectively denoises noisy measurements.

We next applied SPgen to infer proteome-wide spatial distributions for 17,230 mouse proteins from UniProt. Clustering of the predicted spatial profiles revealed two major spatial domains within the intestinal villi. The first domain corresponds to the apical epithelial region of the intestinal villi. This domain is enriched for canonical epithelial and enterocyte markers, including APOA4, APOA1, CDH1, and DPEP1 (**Fig. 2d**). Gene Ontology (GO) analysis of the top 200 differentially expressed (DE) proteins highlights mitochondrial translation, translational elongation, and peptide biosynthetic processes (**Fig. 2e**), consistent with the high metabolic and biosynthetic activity of absorptive enterocytes in the villus apical region. In contrast, the second domain corresponds to the subepithelial region beneath the villus epithelium. This domain is enriched for smooth muscle–associated markers, including CNN1, ACTA2, ACTG2, and MYH11 (**Fig. 2f**). GO analysis reveals strong enrichment of muscle contraction, actin–myosin filament sliding, actomyosin organization, and myofibril assembly pathways (**Fig. 2g**), consistent with the presence of smooth muscle and contractile stromal components underlying the epithelium. Together, these results demonstrate that SPgen recapitulates biologically meaningful proteome-wide spatial organization that reflects known anatomical architecture of the intestinal villi.

### SPgen generalizes to mouse brain spatial proteomics and agrees with HPA measurements

We next evaluated SPgen on a mouse brain coronal spatial proteomics dataset comprising 208 spatial locations and 4,351 measured proteins. Proteins were split into training and testing sets (9:1), as in the previous analysis. SPgen accurately reconstructed spatial protein distributions for held-out proteins, particularly those with strong spatial heterogeneity. Representative examples reveal clear region-specific expression patterns consistent with known cortical–subcortical organization (**Fig. 3a**).

**Figure 3.**
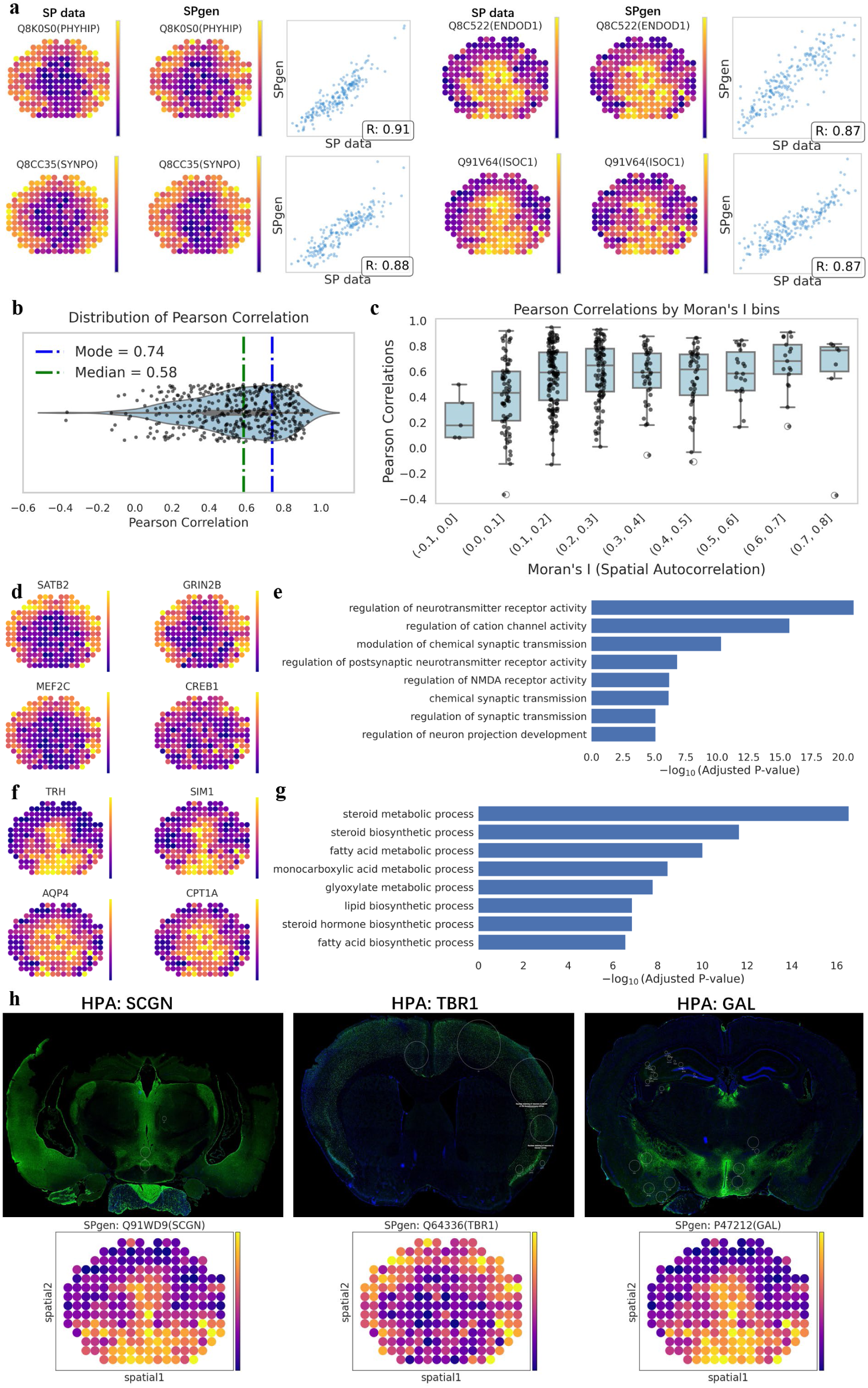
Results on the mouse brain coronal SP data. (a) Representative examples of SPgen performance on the testing set, showing observed and predicted spatial distributions and their joint distribution. Pearson correlation coefficients are indicated. (b) Distribution of Pearson correlation on the testing set, with mode and median annotated. (c) Relationship between testing performance and protein spatial autocorrelation. (d,f) Predicted expression of representative proteins highly expressed in spot clusters 0 (panel d) and 1 (panel f). (e,g) GO enrichment analysis of the top 200 DE proteins in spot clusters 0 (panel e) and 1 (panel g). (h) Comparison between the HPA samples and for SCGN, TBR1 and GAL. HPA samples are downloaded from the mouse brain proteome resource at: https://www.proteinatlas.org/humanproteome/brain/mouse+brain.

Across testing proteins, SPgen achieved a median Pearson correlation of 0.58 (mode 0.73), with performance varying according to the degree of spatial structure in the underlying proteins (**Fig. 3b**). Proteins exhibiting stronger spatial autocorrelation showed substantially higher predictive accuracy, indicating that SPgen preferentially captures biologically structured spatial signals (**Fig. 3c**).

We further applied SPgen to generate proteome-wide spatial profiles for 17,230 mouse proteins. Clustering of predicted spatial patterns revealed two major brain regions. One cluster was enriched in cortical regions and included canonical neuronal markers such as SATB2, GRIN2B, MEF2C, and CREB1, with GO enrichment highlighting synaptic signaling and neurotransmitter receptor regulation (**Fig. 3d,e**). The second cluster was associated with central and ventral brain regions and included hypothalamic markers such as TRH, with enrichment of steroid metabolic and neuroendocrine pathways (**Fig. 3f,g**).

Finally, we benchmarked SPgen predictions against antibody-based spatial protein profiles from the Human Protein Atlas (HPA) mouse brain dataset. As shown in **Fig. 3h**, SPgen recapitulates the spatial patterns of SCGN, TBR1, and GAL in HPA data, despite these proteins being unseen during training. Results for all proteins in the mouse brain spatial proteome of HPA are shown in **Fig. S10**.

### SPgen generalizes to cerebellar spatial proteomics and reveals cross-modal tissue organization

We next applied SPgen to a cerebellar spatial proteomics dataset from PLATO, comprising 1,677 spatial locations and 1,399 proteins. Using the same training–testing split as in previous analyses, SPgen accurately reconstructed spatial protein distributions for held-out proteins (**Fig. 4a**). Representative examples highlight region- specific expression patterns consistent with the annotated cerebellar architecture (**Fig. 4b**).

**Figure 4.**
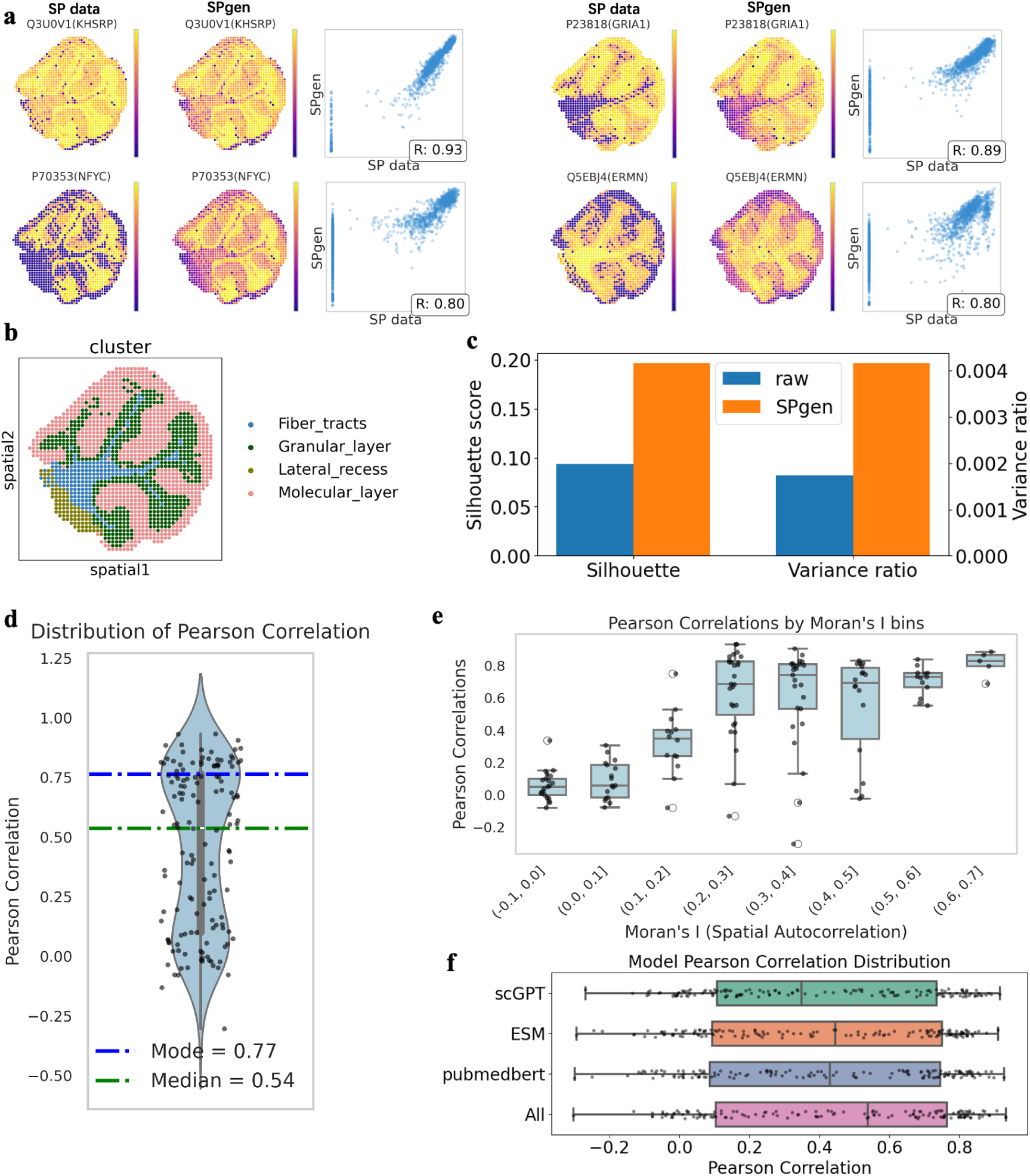
Reconstruction of SP on the Cerebellum dataset. (a) Representative examples of SPgen performance on the testing set, showing observed and predicted spatial distributions and their joint distribution. Pearson correlation coefficients are indicated. (b) The annotation of spatial clusters. (c) Comparison of spatial consistency with annotated clusters between SP data and SPgen predictions. Higher Silhouette scores and variance ratios (between-cluster variance divided by within-cluster variance) indicate stronger agreement with the annotations. (d) Distribution of Pearson correlation on the testing set, with mode and median annotated. (e) Relationship between testing performance and protein spatial autocorrelation. (f) Ablation analysis of protein embeddings used in SPgen.

To evaluate consistency with anatomical structure, we compared spatial organization derived from both measured and predicted protein profiles against annotated cerebellar regions, including fiber tracts, granular layer, lateral recess, and molecular layer. SPgen predictions achieved higher Silhouette scores and between- to within-cluster variance ratios than the original measurements, indicating improved agreement with known anatomical organization (**Fig. 4c**). Across all proteins in the testing set, SPgen achieved a median Pearson correlation of 0.54, with a mode of 0.76 (**Fig. 4d**). Consistent with previous datasets, SPgen’s performance increasing for proteins exhibiting stronger spatial variation (**Fig. 4e**). Ablation analysis demonstrates that the fusion of multi- modality protein embeddings consistently outperforms any individual embedding modality (**Fig. 4f**).

We further generated proteome-wide spatial profiles for 17,230 mouse proteins and identified region-specific protein signatures across cerebellar structures. The molecular layer was enriched for synaptic proteins including VAMP2, GRIA1, GRID2, and SYT5, with GO enrichment highlighting synaptic signaling and GPCR-mediated pathways (**Fig. 5a,b**). The lateral recess exhibited enrichment of transport-related processes consistent with exchange functions at brain interfaces (**Fig. 5c,d**), while fiber tracts showed strong enrichment of myelin-associated proteins including PLP1, MOG, MBP, and MYRF, reflecting compact myelinated axonal bundles (**Fig. 5e,f**). Notably, PLP1 and MOG are membrane-associated myelin proteins that are often underrepresented in mass spectrometry–based spatial proteomics due to the technical challenges. Nevertheless, SPgen consistently reconstructed their spatial distributions, revealing strong enrichment in white matter fiber tracts, in agreement with the abundance of myelinated axonal bundles in these regions.

**Figure 5.**
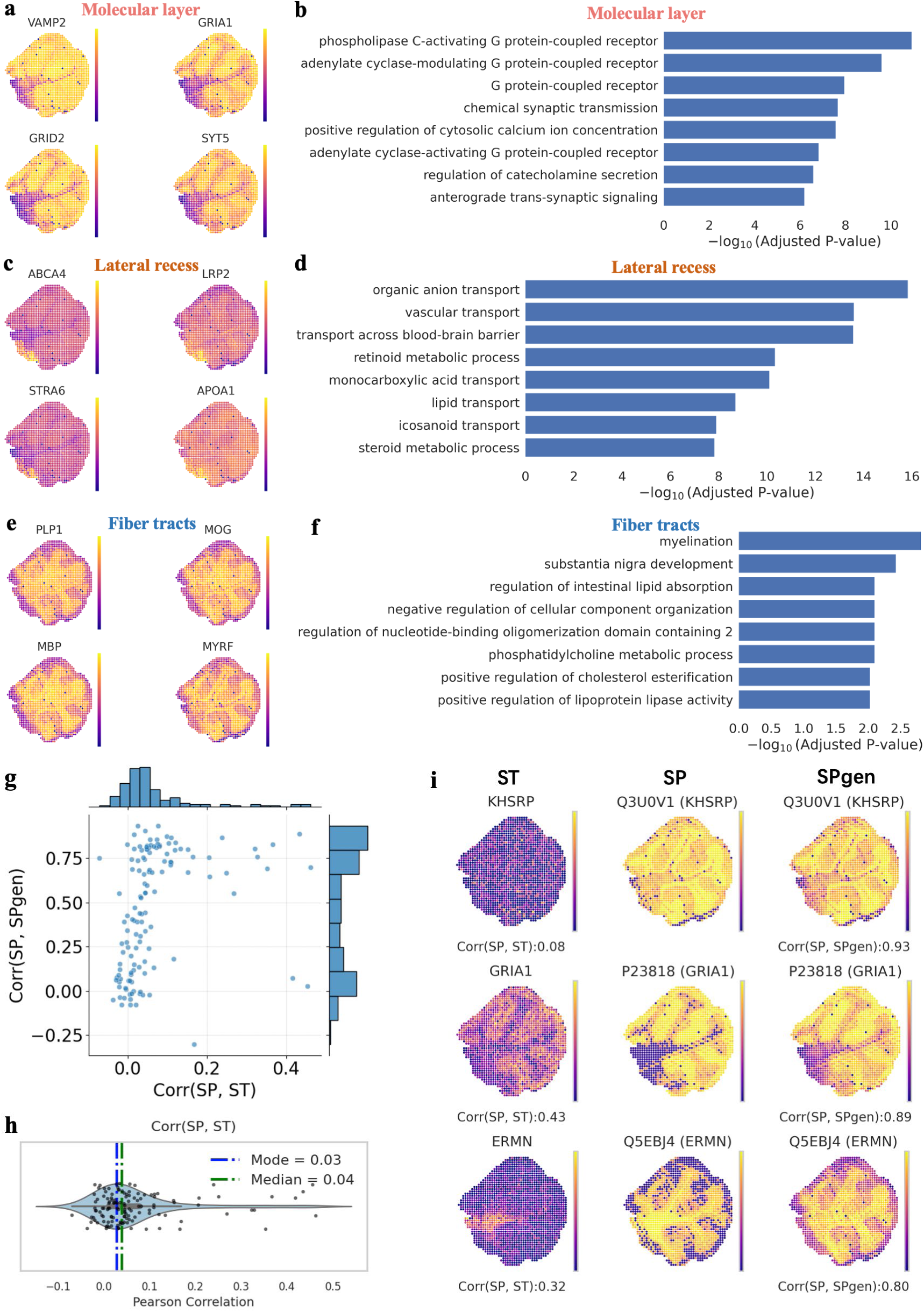
Proteome-wide SP generation and multi-omics analysis on the Cerebellum SP dataset. (a) Predicted expression of representative proteins highly expressed in the molecular layer. (b) GO enrichment analysis of the top 200 DE proteins in the molecular layer. (c) Predicted expression of representative proteins highly expressed in the lateral recess. (d) GO enrichment analysis of the top 200 DE proteins in the lateral recess. (e) Predicted expression of representative proteins highly expressed in the fiber tracts. (f) GO enrichment analysis of the top 200 DE proteins in the fiber tracts. (g) Joint distribution of transcriptome–proteome and SPgen–proteome correlations. (h) Distribution of proteome -transcriptome correlations. (i) The muti-omics analysis for representative proteins. For each protein, the panel displays the corresponding gene’s transcript abundance, the measured protein abundance in the SP data, and the SPgen-predicted protein abundance (from left to right).

Finally, leveraging matched spatial transcriptomics data, we assessed cross-modal consistency between RNA and protein spatial organization^27^. While correlations between spatial transcriptomics (ST) and measured proteomics were generally weak, SPgen predictions showed substantially higher agreement with protein-level spatial patterns. Representative comparisons across ST, measured SP, and SPgen predictions demonstrate that SPgen recovers spatially structured protein distributions that are not directly reflected in transcriptomic profiles (**Fig. 5g–i**), highlighting its ability to capture protein-level spatial organization in complex brain tissues.

## Discussion

Spatial omics is rapidly advancing our ability to resolve tissue organization at molecular resolution. Recent mass spectrometry–based SP technologies enable the quantification of thousands of proteins per spatial location, substantially improving proteome coverage. However, this coverage remains far from complete, and is fundamentally constrained by the high cost and throughput limitations of MS-based assays. As a result, there is an inherent trade-off between proteome coverage and spatial resolution, which limits large-scale, routine profiling and constrains biological discovery in spatial proteomics. These limitations highlight the need for computational approaches that can extend effective protein coverage beyond experimentally measurement.

Here we present SPgen, a multi-modal framework for proteome-wide spatial protein prediction. SPgen integrates protein sequences, functional annotations, and genomics signals to learn transferable representations across molecular modalities. Across multiple SP datasets, SPgen accurately reconstructs spatial patterns of unseen proteins, enabling proteome-scale spatial prediction and supporting downstream analyses. In addition, SPgen improves signal consistency in measured data by reducing spatial noise, thereby enhancing the biological interpretability of SP profiles.

Despite these advances, SPgen is currently trained and evaluated on individual SP datasets, which may limit its generalization across platforms. This is partly due to the intrinsic noise and sparsity of MS-based spatial proteomics and the limited and imbalanced coverage of protein measurements used for supervision, which may introduce dataset-specific biases.

Looking forward, SPgen provides a general framework for scalable spatial proteomics modeling. Its modular design allows straightforward integration of additional modalities, such as spatial transcriptomics and imaging-based measurements. With the continued expansion of spatial multi-omics technologies, SPgen may serve as a foundation for cross-platform models capable of predicting spatial protein distributions across tissues and experimental conditions.

## Methods

### Preprocessing for SP data

Raw protein count matrices (*Y_rarr_*) for both the training and testing subsets are firstly log-transformed and standardized on a per-gene basis. For each dataset, expression values are z-score normalized by subtracting the mean and dividing by the standard deviation across spatial spots:

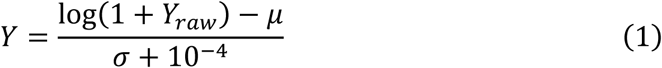

where *μ* and *σ* denote the gene-wise mean and standard deviation, respectively, and a small constant (1e-4) is added to the denominator to avoid numerical instability. The normalized expression matrices are used for all subsequent model inputs.

For each spatial proteomics dataset, proteins are randomly partitioned into training and testing sets at a 9:1 ratio, and the normalized training matrix is used as the primary model input.

Spatial dependencies among tissue locations are characterized using spot coordinates embedded in the original Visium array. A spatial graph is constructed by connecting each spot to its four nearest neighbors (K=4) based on Euclidean distance, resulting in an undirected adjacency structure used for downstream spatial inference.

To quantify gene-level spatial structure, Moran’s I statistics are computed independently for the training and test sets using the python package squidpy. For each dataset, spatial neighbor graphs are first computed (sq.gr.spatial_neighbors), and Moran’s I values are estimated using a permutation-based test (sq.gr.spatial_autocorr, 1000 permutations). The resulting Moran’s I scores are extracted for all profiled proteins and used to interpret spatial predictability in benchmarking analyses.

### Protein embedding learned from multi-modality foundation models

We integrate three complementary protein representations to capture distinct molecular characteristics, which are textual, and amino-acid–sequence embeddings and co-expression.

To leverage the rich transcriptional information derived from high-throughput single-cell datasets, we adopt co-expression embeddings generated by the pre-trained scGPT model (‘‘whole-human’’ mode), which is trained on 33 million cells and recommended by its author (https://github.com/bowang-lab/scGPT). Gene-level embeddings are mapped to their corresponding proteins using UniProt annotations.

For texture embeddings, we extract descriptive text associated with each protein from UniProt and encoded these annotations using a PubMedBERT model pre-trained on biomedical literature, thereby incorporating curated functional and localization knowledge.

Amino-acid–sequence embeddings are obtained by processing each protein’s primary sequence (retrieved from UniProt) through the pre-trained ESM model, capturing physicochemical and structural determinants encoded at the sequence level.

Together, these three modalities provide a diverse and complementary representation of protein biology. For each protein *i*, we denote its modality-specific embeddings as:

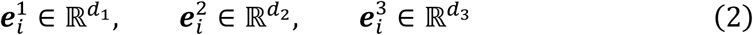

For each modality, a dedicated linear projection layer maps the modality-specific gene representation into a shared latent space of dimension

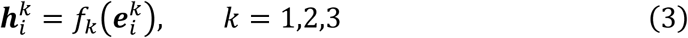

During training, random element-wise masking is applied independently to each modality-specific embedding with masking ratio p, serving as a feature-level regularization strategy. The final fused protein representation is computed as:

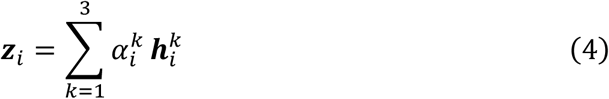

Where 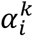 represents modality-specific weights for each gene.

### Learning spot embeddings with GAT

The spot embeddings are learned from spatial proteomics data using a multi-layer GAT. The spatial graph is constructed based on tissue coordinates using a K-nearest neighbors strategy (K = 4), ensuring that each spot is connected to its four closest spatial neighbors. Using the preprocessed protein abundance vectors from the training set as node features, the GAT model propagates local spatial information through four stacked GATConv layers, each followed by layer normalization and residual connections to stabilize training and enhance representation capacity.

For the *l* − *t*ℎ layer, suppose X is the node features and *E* is the set of edges, we compute:

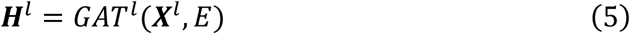

Each GAT layer uses two attention heads and concatenation disabled, which corresponds to averaging the attended outputs. This design enables the learned spot embeddings to capture both local neighborhood structure and broader tissue-level organization.

### The generation of SP profile by SPgen

Suppose the number of proteins and spots are *G* and *N* respectively, after obtaining the protein embeddings *Z* = [*z*_1_, *z*_2_ … *z_M_*] ∈ ℝ*^G^*^∗*d*^ from the fused multi-modality foundation models and the spot embeddings *H* = [ℎ_1_, ℎ_2_ … ℎ*_N_*] ∈ ℝ*^N^*^∗*d*^ from the GAT module, SPgen generates the SP matrix *Y* ∈ ℝ*^N^*^∗*G*^ through a bilinear interaction between the two embedding spaces by

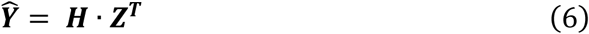

In detail, for each spot–protein pair (*i*, *j*), the generated intensity is: *Ŷ_i_,_j_* =< ℎ*_i_* , *z_j_* >, where ℎ*_i_* is the embedding of spot *i* and *z_j_* is the embedding of protein *j*. This inner-product–based generation could offer several advantages, including computational efficiency, high interpretability and strong generalization with reduced overfitting.

### Training details of SPgen

For those proteins in the training set, suppose their SP profile is *Y*. The loss function is defined as a combination of L1 and L2 reconstruction losses. In addition, recognizing that a subset of proteins is weakly associated with spatial structure, Moran’s I is used as a weighting term in the loss function. This design biases optimization toward spatially variable proteins, ensuring that features carrying meaningful spatial signals have greater influence on model learning. Overall, the loss function is:

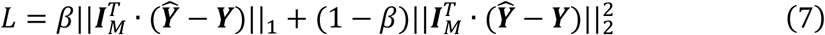

where *β* is a hyper-parameter that set to 0.5, *I_M_* is the vector filled by the Moran’s I scores of all proteins. For each dataset, SPgen is trained on the training set for 8,000 epochs with a learning rate of 0.001. All hyperparameters and optimization settings are held constant across datasets.

### Spatial autocorrelation metrics

To quantify the spatial continuity of expression for each protein, we compute two widely used spatial autocorrelation statistics, Moran’s I and Geary’s C, based on a spatial neighbor graph.

For a given protein, let *s_i_* denote its expression value at spatial location *i*, with *n* total locations. A spatial weight matrix *M* = (*m_i_*_,*j*_) is constructed using a *k*-Nearest Neighbor (kNN) graph (default *k* = 6), where *w_i_*_,*j*_ > 0 indicates that locations *i* and *j* are spatial neighbors. Locations with no neighbors (isolated nodes) are excluded from the analysis.

We first centered the expression values:

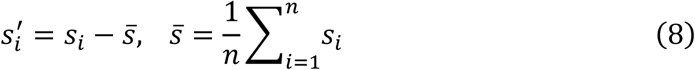

1. Moran’s I

Moran’s I measures global spatial autocorrelation by quantifying the similarity of expression values between neighboring locations:

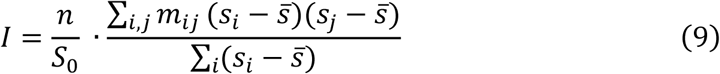

where *S*_0_ = ∑*_i_*_,*j*_ *m_ij_* is the total sum of spatial weights. Higher values of Moran’s I indicate stronger positive spatial autocorrelation (i.e., neighboring locations tend to have similar expression levels), while values near zero indicate spatial randomness.

(2) Geary’s C

Geary’s C measures local spatial dissimilarity by directly comparing differences between neighboring locations:

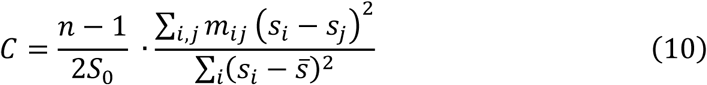

Unlike Moran’s I, lower values of Geary’s C indicate stronger spatial autocorrelation. A value of *C* = 1 corresponds to spatial randomness, *C* < 1 indicates positive spatial autocorrelation, and *C* > 1 indicates negative spatial autocorrelation.

### Evaluation with Pearson correlation

On each SP dataset, after being trained on the training set comprising thousands of proteins, SPgen is capable of generating spatial distributions for any queried protein on the corresponding tissue, based on its embedding from the multi-modality foundation model. To evaluate predictive accuracy, we first inferred spatial maps for proteins in the testing partition. The Pearson correlation between predicted and experimentally measured expression patterns is employed as the evaluation metric. The joint distribution of testing scores and spatial autocorrelation (Moran’s I) enables assessment of how SPgen performance varies with the underlying spatial structure of individual proteins.

### The proteins clustering

To stratify proteins according to their spatial expression patterns, we performed unsupervised clustering based on their spot-level abundance profiles. We standardized each protein expression vector to zero mean and unit variance using z-score scaling, ensuring that differences in magnitude did not dominate downstream analyses. Principal component analysis (PCA) is then applied to obtain a low-dimensional representation of protein-level variation. PCA is performed using 50 components and a deterministic random seed at 42 to ensure reproducibility. The resulting principal components are subsequently subjected to k-means clustering, grouping proteins into clusters according to their distribution patterns.

### The spots clustering and the large-scale spatial protein analysis

To delineate spatially coherent tissue domains, we applied unsupervised clustering to spot-level protein expression profiles. We first selected the top 2000 spatially variable proteins according to the Moran’s I, and perform principal component analysis to retain the top 50 components to capture the dominant axes of variation across spots. A shared nearest-neighbor graph is subsequently constructed in the reduced space using 15 neighbors and the first 15 principal components. Spatial clusters are then inferred from this graph using the Leiden community detection algorithm. This procedure robustly recovers putative tissue domains and uncovers mesoscale organization in protein expression that is broadly consistent with histological structure.

To enable large-scale spatial protein profiling, we aggregate all proteins represented across the three embedding spaces, yielding a catalogue of 573,230 unique protein entries. Among these, 17,230 proteins are annotated as Mus musculus. Differential expression analysis is performed independently for each spatial cluster based on the generated spatial profiles.

Gene Ontology (GO) enrichment analysis is conducted using the gseapy Python package. The top 200 cluster-specific proteins ranked by differential expression are used as input, and the full set of 17,230 Mus musculus proteins serve as the background universe. Significantly enriched biological processes are identified based on adjusted p-values.

### The ablation study on multi-modality protein embeddings

We perform an ablation analysis to assess the contribution of each protein embedding modality to the overall model performance. Specifically, we considered three embedding sources: (i) expression-informed representations with scGPT, (ii) sequence-derived representations from ESM, and (iii) text-informed representations with pubmedBERT. For each modality, a separate model variant is trained by disabling one or more embedding inputs while keeping all other components and training hyperparameters fixed.

The ablations are conducted under identical training and testing sets within each spatial proteomics dataset. Model performance is evaluated using the same metrics used in the baseline setting, enabling direct comparison among single-modality, dual-modality, and full multi-modality configurations.

## Data Availability Statement

The protein annotations, including amino acid sequences and functional descriptions, are obtained from UniProt at https://www.uniprot.org/help/downloads.

Three pretrained foundation models are employed in this study, including the scGPT model from https://github.com/bowang-lab/scGPT/, the ESM model from https://dl.fbaipublicfiles.com/fair-esm/models/esm1b_t33_650M_UR50S.pt, and the pubmedBERT model from https://huggingface.co/NeuML/pubmedbert-base-embeddings.

Three SP datasets used in this study are released by previous studies. The intestinal villi dataset and Cerebellum dataset are released by PLATO study, available at https://github.com/bioinfo-biols/Flow2Spatial/tree/main/tests/adata.h5ad and https://github.com/bioinfo-biols/Flow2Spatial/tree/main/datasets respectively. The mouse brain coronal dataset is released by Spatial-DC study, available at https://zenodo.org/records/14523511.

## Code Availability Statement

TSvelo is implemented in Python. The source code can be downloaded from the GitHub repository, https://github.com/lijc0804/SPgen.

## Acknowledgement

This work is supported by National Natural Science Foundation of China (62503452).

## Competing Interests Statement

The authors declare no competing interests.

## Notes

### Competing Interest Statement

The authors have declared no competing interest.

